# HypercubeME: two hundred million combinatorially complete datasets from a single experiment

**DOI:** 10.1101/741827

**Authors:** Laura Avino Esteban, Lyubov R. Lonishin, Daniil Bobrovskiy, Gregory Leleytner, Natalya S. Bogatyreva, Fyodor A. Kondrashov, Dmitry N. Ivankov

## Abstract

**Motivation:** Epistasis, the context-dependence of the contribution of an amino acid substitution to fitness, is common in evolution. To detect epistasis, fitness must be measured for at least four genotypes: the reference genotype, two different single mutants and a double mutant with both of the single mutations. For higher-order epistasis of the order n, fitness has to be measured for all 2^n^ genotypes of an n-dimensional hypercube in genotype space forming a “combinatorially complete dataset”. So far, only a handful of such datasets have been produced by manual curation. Concurrently, random mutagenesis experiments have produced measurements of fitness and other phenotypes in a high-throughput manner, potentially containing a number of combinatorially complete datasets.

**Results:** We present an effective recursive algorithm for finding all hypercube structures in random mutagenesis experimental data. To test the algorithm, we applied it to the data from a recent HIS3 protein dataset and found all 199,847,053 unique combinatorially complete genotype combinations of dimensionality ranging from two to twelve. The algorithm may be useful for researchers looking for higher-order epistasis in their high-throughput experimental data.

**Availability:** https://github.com/ivankovlab/HypercubeME.git.

## Introduction

Epistasis, the dependence of the impact of a mutation on the genetic context, is abundant and important phenomenon in molecular evolution [1]. Formally, epistasis is characterized by coefficients α having two or more indices in the following representation of fitness f as a function of a genotype g (assuming, for simplicity, that maximum of one mutation is allowed at any position):

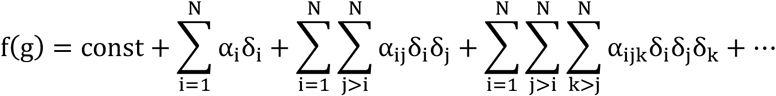

where δ_i_ = 1 if i-th position is mutated in genotype g; otherwise δ_i_ = 0 or δ_i_ = −1, depending on the formalism of epistasis description [2]. Coefficients α_i_ correspond to a single effect of the mutation in the i-th position. Coefficients α_ij_, having two indices, represent the minimal possible pair-wise epistasis between positions i and j, while coefficients α having three or more indices correspond to “higher-order epistasis” [2-9].

As can be seen from the equation above, to detect epistasis of the n-th order, one has to measure phenotypes of all 2^n^ genotypes forming an n-dimensional hypercube in genotype space. Such experimental datasets are called “combinatorially complete datasets” [3]. Up to now, higher-order epistasis was studied using only a handful of examples carefully designed to have all 2^n^ combinations [3]. On the other hand, high-throughput experiments using (quasi-)random mutagenesis have produced vast amounts of data: 51,715 measured genotypes for GFP [10], over 65,000 for arginine tRNA [11], 956,648 for HIS3 protein [12], to name a few. These experiments may contain a number of combinatorially complete datasets as subsets of a general dataset. However, the extraction of such hypercubes from a large dataset is not straightforward, and may not be feasible to do in a brute-force manner.

We suggest here a simple recursive algorithm to extract all combinatorially complete datasets from a high-throughput (quasi-)random mutagenesis dataset. To illustrate it, we applied it to the recently published data on the mutagenesis of HIS3, the largest experimentally measured dataset to date, containing fitness measurements for about one million genotypes [12]. We have identified about 200 million combinatorially complete datasets, of dimensionality from two to twelve. Squares (two-dimensional hypercubes) comprise 12% of all hypercubes, while the rest 88% provide information about higher-order epistasis.

## Algorithm

The algorithm uses the fact that any n-dimensional hypercube contains two opposite hyperfacets, which are parallel to each other. Those hyperfacets are hypercubes of dimensionality (n − 1), which, in turn, are built from parallel hypercubes of dimensionality (n − 2), etc., down to hypercubes of 0-th dimensionality (which are simply points in genotype space, i.e., genotypes).

The algorithm consists of repeating steps. At each step, all possible n-dimensional hypercubes are generated from the set of (n − 1)-dimensional hypercubes. Informally, at each step, the algorithm takes all pairs of existing parallel hypercubes and if the distance between the hypercubes in the pair is one, the pair composes the hypercube of a higher dimensionality (Fig.1). We need to define the diagonal of a combinatorially complete dataset as a list of mutations transforming a genotype of the dataset to the most distant genotype of the same dataset. An n-dimensional combinatorially complete dataset therefore contains 2^n − 1^ diagonals, each of which (if not empty) can be written in the forward and reverse direction.

**Figure 1.**
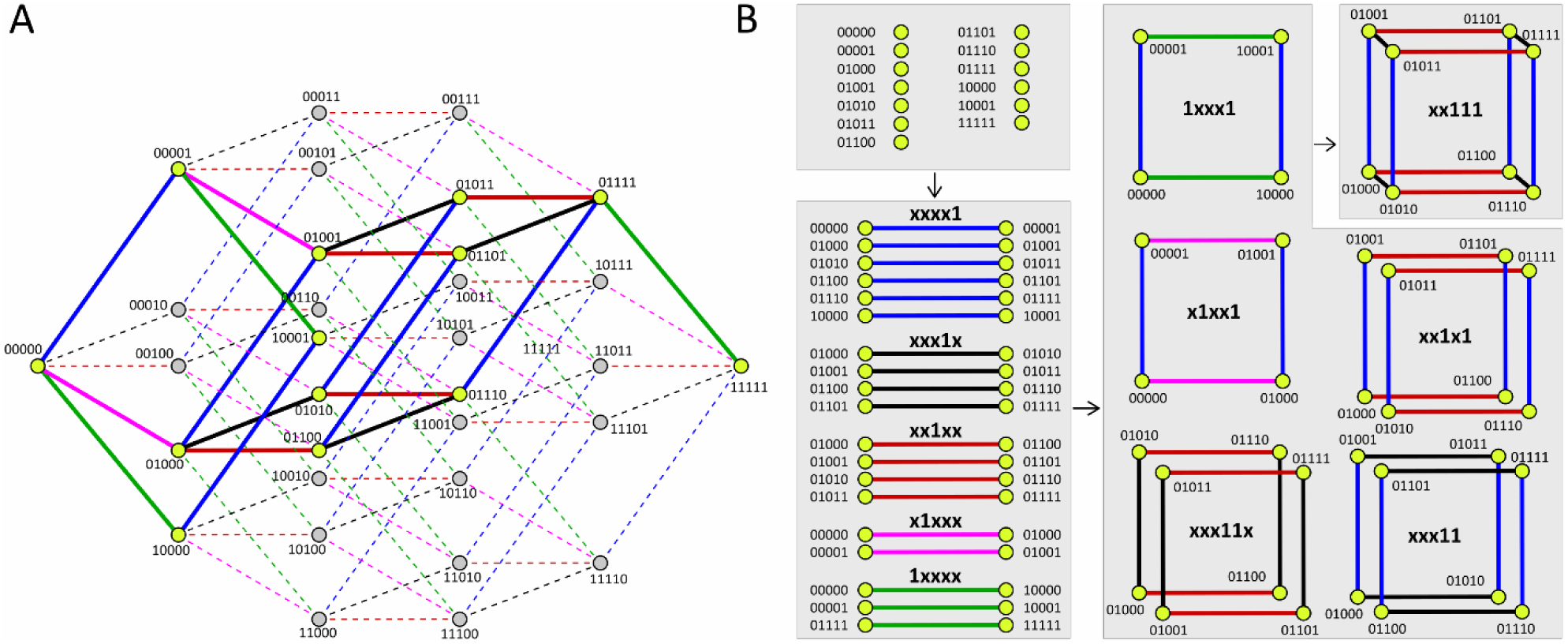
Example illustrating the work of the algorithm. (A) A graph corresponding to a five-dimensional hypercube is given, where measured genotypes are drawn in yellow-green and non-measured genotypes are drawn in grey. Vertices are given in binary notation, where each digit corresponds to one of five substitution sites. The digit is zero if the corresponding substitution site contains an amino acid of the reference ‘00000’ genotype; otherwise, it is one. Two vertices are connected if they differ by only one digit. The graph is drawn so that: (1) all vertices are visible (that is, no vertex is shaded by another one); (2) all vertices having the same number of zeros belong to the same vertical line; (3) the edges parallel in 5-dimensional cube are drawn parallel to each other, and in different colors, for convenience. Blue, black, red, pink, and green edges correspond to substitutions in the fifth, fourth, third, second, and first sites, respectively. (B) The algorithm is applied to an example of random mutagenesis data from the panel (A). The diagonal for each group of hypercubes is given in bold.

Formally, the algorithm consists of the following steps:

1. **Input data.** The algorithm takes the list of genotypes as input data. Each genotype is a point in genotype space, being a hypercube of dimensionality zero. The diagonal of each such hypercube is zero.
2. **Construction of 1-dimensional hypercubes from input data.** The algorithm builds all genotype pairs at distance of a single amino acid substitution: it goes through all genotype pairs and remembers those pairs in which genotypes differ by one mutation. These found pairs are hypercubes of dimensionality one with the diagonal being the difference between the two genotypes in a pair. The pair is stored in such an order that the diagonal is alphabetically less than the alternative variant.
3. **Construction of (n + 1)-dimensional hypercubes from n-dimensional ones.** The hypercubes of dimensionality n produced in the previous step are divided into groups having the same diagonal. They are parallel to each other in genotype space. In each group, the algorithm goes through all pairs of n-dimensional hypercubes, and if the distance between two hypercubes in a pair is one, the combination of the two hypercubes results in an (n + 1)-dimensional hypercube. The diagonal of the found hypercube is formed by concatenating the diagonal of the two n-dimensional hypercubes with the mutation between them. Again, the direction of the mutation is chosen to be alphabetically less than for the alternative one. To avoid duplicates, the found (n + 1)-dimensional hypercube is stored only if the mutations in the diagonal are ordered alphabetically. This step is repeated until no new hypercubes are found.

Each step of the algorithm can be easily parallelized. The multi-processor version can be found at https://github.com/ivankovlab/HypercubeME.git.

## Results

We have applied the algorithm to the recently published fitness landscape for HIS3 protein [12], the biggest fitness landscape published so far. The HIS3 protein was divided into twelve segments, and quasirandom mutagenesis has been done in each segment separately. We had to exclude indels and mutations outside the segment, so the number of considered experimentally measured genotypes ranged from 16,182 for segment S7 to 82,081 for segment S2, overall summing up to 721,791 genotypes (Table S1).

We have found all 199,847,053 hypercubes having dimensionality from two to twelve. The single-processor working time ranged from two hours for S7 to almost ten days for S5. Among the found hypercubes, the percentage of squares was 12%, while the remaining 88% had dimensionality 3 and higher and, thus, can be used for exploring higher-order epistasis. The number of found hypercubes throughout segments is given in Table S2.

## Acknowledgements

We are grateful to Cathy Shufro for help with editing the text.

