## Supplemental Table 1 for "HypercubeME: two hundred million combinatorially complete datasets from a single experiment"

Table S1. Number of measured genotypes

| Segment | S1 | S2 | S3 | S4 | S5 | S6 | S7 | S8 | S9 | S10 | S11 | S12 | All |
|---|---|---|---|---|---|---|---|---|---|---|---|---|---|
| Number of genotypes | 60 757 | 82 081 | 68 867 | 63 842 | 72 801 | 64 187 | 16 182 | 59 453 | 78 576 | 63 238 | 34 478 | 57 329 | 721 791 |
