## Supplemental Table 2 for "HypercubeME: two hundred million combinatorially complete datasets from a single experiment"

### Table S2. Number of hypercubes found in HIS3 experiment

| Dimension / Segment | 2 | 3 | 4 | 5 | 6 | 7 | 8 | 9 | 10 | 11 | 12 | All |
|---|---|---|---|---|---|---|---|---|---|---|---|---|
| S1 | 3 239 975 | 8 047 443 | 11 488 387 | 9 671 672 | 4 731 655 | 1 266 830 | 161 069 | 6 586 | – | – | – | 38 613 617 |
| S2 | 2 748 491 | 5 577 678 | 6 592 980 | 4 677 409 | 1 985 551 | 487 450 | 63 938 | 3 679 | 50 | – | – | 22 137 226 |
| S3 | 2 425 273 | 5 369 736 | 7 177 755 | 5 940 945 | 3 004 618 | 881 893 | 134 673 | 8 399 | 91 | – | – | 24 943 383 |
| S4 | 1 721 693 | 3 161 825 | 3 464 103 | 2 323 212 | 940 378 | 219 068 | 26 433 | 1 248 | 9 | – | – | 11 857 969 |
| S5 | 2 116 191 | 4 546 950 | 6 289 934 | 5 901 892 | 3 855 482 | 1 764 696 | 560 241 | 120 337 | 16 549 | 1 257 | 33 | 25 173 562 |
| S6 | 1 646 629 | 2 967 579 | 3 100 174 | 1 925 900 | 705 769 | 148 593 | 17 204 | 1 007 | 20 | – | – | 10 512 875 |
| S7 | 748 123 | 1 853 730 | 2 578 331 | 1 977 559 | 758 004 | 110 924 | – | – | – | – | – | 8 026 671 |
| S8 | 1 471 038 | 2 425 504 | 2 247 568 | 1 186 009 | 343 808 | 49 991 | 2980 | 34 | – | – | – | 7 726 932 |
| S9 | 1 427 354 | 1 734 410 | 1 024 513 | 273 748 | 28 859 | 896 | 3 | – | – | – | – | 4 489 783 |
| S10 | 2 515 012 | 5 565 341 | 6 988 005 | 4 999 483 | 1 967 171 | 402 753 | 39 792 | 1 454 | – | – | – | 22 479 011 |
| S11 | 1 425 638 | 3 079 322 | 3 593 920 | 2 173 218 | 598 358 | 54 398 | 381 | – | – | – | – | 10 925 235 |
| S12 | 1 789 695 | 3 475 957 | 3 897 708 | 2 576 913 | 988 823 | 209 135 | 21 741 | 817 | – | – | – | 12 960 789 |
| All | 23 275 112 | 47 805 475 | 58 443 378 | 43 627 960 | 19 908 476 | 5 596 627 | 1 028 455 | 143 561 | 16 719 | 1 257 | 33 | 199 847 053 |
